# SP1433-1438 operon of *Streptococcus pneumoniae* regulates metal homeostasis and cellular metabolism during zinc-stress

**DOI:** 10.1101/367086

**Authors:** Lindsey R. Burcham, Rebecca A. Hill, Rachel C. Caulkins, Joseph P. Emerson, Bindu Nanduri, Jason W. Rosch, Nicholas C. Fitzkee, Justin A. Thornton

**Author notes:** Corresponding author: Justin A. Thornton.

## Abstract

*Streptococcus pneumoniae* colonizes the mucosa of the human nasopharynx and is a leading cause of community-acquired pneumonia, acute otitis media, and bacterial meningitis. Metal ion homeostasis is vital to the survival of this pathogen and contributes significantly to both colonization and invasive disease. Microarray and qRT-PCR analysis revealed an upregulation of an uncharacterized operon (*SP1433-1438*) in pneumococci subjected to metal-chelation by *N,N,N’,N’*-tetrakis-(2-Pyridylmethyl)ethylenediamine (TPEN). Supplementation of either zinc or cobalt following TPEN treatment drastically abrogated induction. BLAST analysis predicted this operon to encode two ABC-transporters, sharing homology to a multidrug resistance system (SP1434-1435) and an energy-coupling factor (ECF) transport system (SP1436-1438). Inductively coupled plasma mass spectrometry (ICP-MS) analysis indicated changes in intracellular concentrations of iron, zinc, and manganese ions in a Δ1434-8 strain compared to parental T4R. Analysis of the secreted metabolomic profile of the T4R and Δ1434-8 strains identified significant changes in pneumococcal glycolytic pathways, indicating a shift towards increased production of acetate. Additionally, proteomic analysis revealed 41 differentially expressed proteins in the Δ1434-8 strain, with roughly 20% of them regulated by the global catabolite repressor, CcpA. Based on these findings, we propose that the *SP1433-1438* operon is largely involved in the central metabolism of *S. pneumoniae* during zinc-limitation.

**Importance:** Metal sequestration is a common strategy utilized by the host immune response as well as antibiotics such as vancomycin to kill invading bacterial pathogens (1). However, pneumococcus is still able to thrive under zinc-limiting conditions. This study describes a previously uncharacterized operon encoding two ABC transport systems that are strongly induced during zinc-limiting conditions. This operon was found to be regulated by a zinc-dependent regulator (*SP1433*) that functions independently of the overarching AdcR regulon. We have additionally utilized a 2D-NMR approach to analyze the secreted metabolome and have employed proteomic analysis to identify a role for these systems in the maintenance of cellular metabolism. This study provides new information on how *Streptococcus pneumoniae* responds and adapts to zinc-limiting conditions.

## Introduction

Bacteria have evolved a wide variety of ATP-binding cassette (ABC) transporters that function primarily to transport molecules across cell membranes (2). These systems are involved in the uptake and efflux of many substrates, including vitamin B12, iron-binding siderophores, and free metal ions (3-5). These transport systems consist of importers, found only in prokaryotic systems (types I, II, and III), and exporters, found in both prokaryotic and eukaryotic systems (2). Type III importers, or energy coupling factor (ECF) transporters, were the most recently characterized, and they differ from importer types I and II in that they lack substrate-binding domains or proteins (6). ECF transporters are involved in the uptake of vitamins (thiamine and riboflavin) and metal ions (cobalt and nickel) (7-9).

In an effort to starve bacterial pathogens of essential metals, the human immune system expresses proteins that sequester metal ions, a process termed “nutritional immunity” (10). As organisms continue to evade the immune response and evolve resistance to antibiotics, metal homeostasis is an attractive target for future therapeutics. Discerning the mechanisms by which pathogens respond to and overcome metal starvation is key to understanding the physiology of the organism and developing novel therapeutics to eliminate them.

*Streptococcus pneumoniae*, pneumococcus, is a Gram-positive commensal of the human nasopharynx and is the leading cause of community acquired pneumonia worldwide (11). Zinc transport systems and the zinc-dependent AdcR regulon of the pneumococcus have been characterized in detail (12-15). However, it remains unknown if other zinc-sensitive transporters exist. Previous work from our laboratory has identified roles for zinc homeostasis in both invasion and biofilm formation of *S. pneumoniae* (12, 16). Here we identify zinc as an effector molecule for the regulation of a genetic locus involved in maintenance of metal homeostasis. We hypothesize that this locus encodes an operon that is zinc-sensitive and largely contributes to cellular metabolism, specifically relating to oxidative stress, carbohydrate metabolism, and metal ion uptake. Additionally, this study has utilized 2D NMR metabolomics and proteomic analysis to identify metabolic pathways of *Streptococcus pneumoniae* that contribute to homeostasis during zinc-stress.

## Results

### Prediction of Two Uncharacterized ABC-Transporters in Streptococcus pneumoniae

Microarray data of *Streptococcus pneumoniae* strain TIGR4 exposed to the zinc-chelator *N*, *N*, *N*’, *N*’-tetrakis-(2-pyridylmethyl)ethylenediamine (TPEN) identified the gene loci *SP1434-1438* as some of the most highly upregulated genes in response to zinc-limitation (Supplemental Table 1). Due to their proximity to each other in the genome (Fig. 1A), and their similar response to zinc-chelation, we hypothesized that these genes comprise an operon involved in homeostasis during zinc-limitation. Analysis using the Database of prOkaryotic OpeRons (DOOR) supported our hypothesis that these genes are located within an operon (17). Co-regulation of genes *SP1434-1438* was verified by detecting a significant upregulation of each gene by quantitative real time PCR (qRT-PCR) following zinc-chelation with TPEN (data not shown). InterPro analysis identified SP1434 and SP1435 as ABC transporter ATP-binding proteins, and BLAST analysis identified both SP1434 and SP1435 as multidrug resistance-like ATP-binding proteins (mdlB) (Supplemental Table 2). Though SP1434 and SP1435 were proposed to be involved in antibiotic efflux (18), when tested against a broad panel of antibiotics, a mutant strain lacking *SP1434* was equally as sensitive as the parental T4R (Supplemental Figure 1). SP1434 (orange) and SP1435 (red) are predicted to form an independent system with each protein containing five alpha helix transmembrane domains and a P-loop ATPase domain (Fig. 1B). Additionally, InterPro analysis indicated SP1436 as a conserved integral membrane protein, SP1437 as a membrane protein, and SP1438 as a cobalt(II) ABC transporter permease. BLAST analysis indicated that SP1436, SP1437, and SP1438 share sequence homology with components of an energy coupling factor (ECF) transport system, representing the substrate-specific component (EcfS), the transmembrane transporter component (EcfT), and the ATP-binding protein (EcfA2), respectively. The predicted ECF transport system is also shown, whereby, SP1436 (purple) binds the substrate, SP1437 (green) functions as a permease, and SP1438 (blue) acts as an ATPase utilizing energy generated by ATP hydrolysis to import the substrate into the cell.

**Figure 1.**
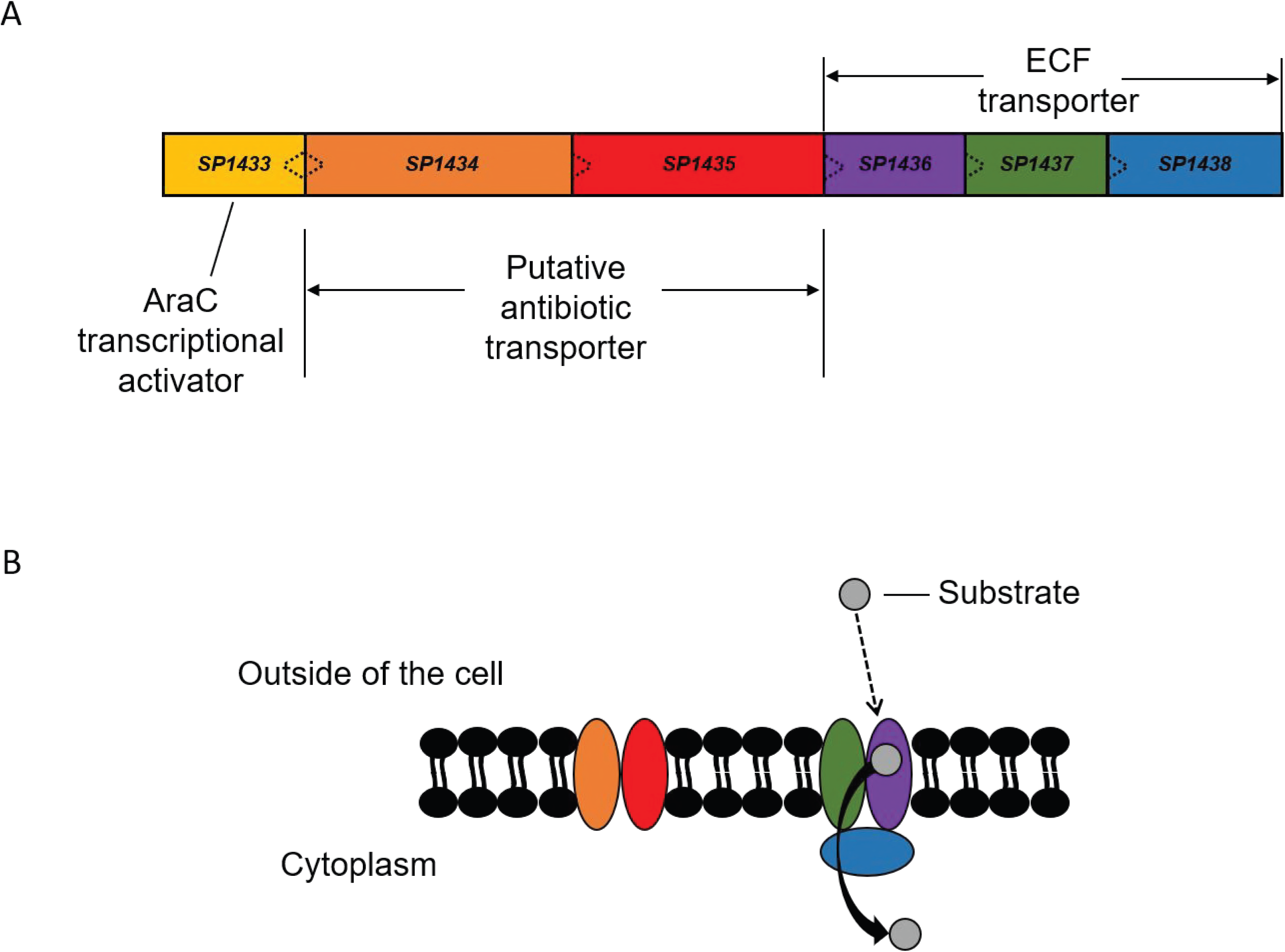
*SP1434-1438* Operon Model. A) the genetic loci containing genes *SP1433-SP1438* with the direction of transcription represented by the dotted arrow. Upstream of this operon is an AraC transcriptional regulator, SP1433 (yellow). B) The working model of the transporters encoded by genes *SP1434* (orange)*-SP1435* (red) and *SP1436-SP1438* (purple, green, and blue, respectively).

#### Metal Availability Alters Expression of SP1434

To verify the microarray results, expression of the first gene in the operon (*SP1434*) was analyzed by qRT-PCR following TPEN treatment. Results from the qRT-PCR analysis indicated robust expression in response to the zinc(II) chelation, as *SP1434* was upregulated greater than 100-fold in comparison to a control sample without TPEN treatment (Fig. 2). In contrast, expression of *SP1434* was unaltered in samples exposed to excess metal ion concentrations; addition of zinc ions led to a 0.7-fold change and adding cobalt(II) (−1.3), iron(II) (−1.1), and nickel(II) (−1.1) yielded similar results. To determine individual metals’ effect on chelation-dependent operon expression, samples were treated with TPEN for 15 min followed by supplementation with excess metal for 15 min. Surprisingly, addition of zinc or cobalt ions limited the upregulation of *SP1434* by roughly 90%, and nickel(II) limited expression by 30% compared to a control treated only with TPEN for 30 min. However, addition of iron(II) following TPEN treatment limited upregulation by less than 10%. This is important to note as TPEN, in addition to binding zinc(II) at a 1:1 ratio (*K*_d_ = 2.6 × 10^−16^ M), also has an affinity for iron(II) (*K*_d_ = 2.4 × 10^−15^ M) (19). Collectively, these data indicate this system is responsive to multiple divalent metal ions, but it does not appear to be affected by iron availability.

**Figure 2.**
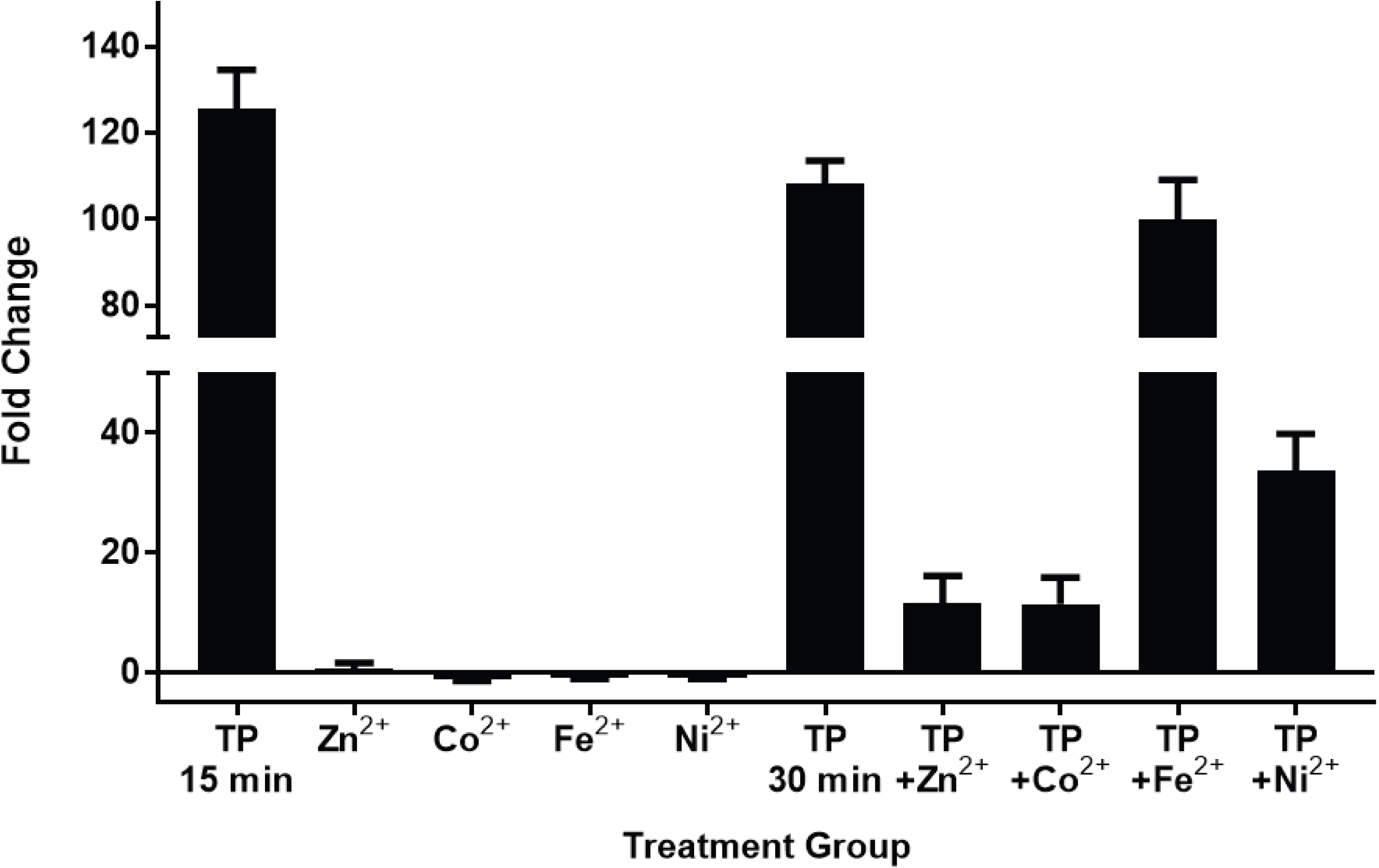
*SP1434* is highly zinc(II) sensitive. Expression of *SP1434* measured by qRT-PCR from RNA extracted following either 15 min treatment with zinc-chelator TPEN (30 μM)/metals (200 μM) or 15 min TPEN (30 μM) exposure with additional 15 min metal supplementation. Fold changes were calculated by ΔΔCT analysis with *gyrA* serving as internal control.

#### Transcriptional Regulation of SP1434-1438 Operon

The locus directly upstream of *SP1434* encodes a previously uncharacterized AraC transcriptional regulator (*SP1433*). To determine if *SP1433* is involved in the induction of *SP1434* following TPEN treatment, expression of *SP1433* and *SP1434* were assessed by qRT-PCR in the parental T4R and a Δ1433 mutant strain. Expression of *SP1433* in the T4R strain following TPEN treatment revealed an upregulation of ≈7 fold (Fig. 3). As expected expression of *SP1433* was undetectable in the Δ1433 mutant. Though *SP1434* was strongly upregulated following TPEN exposure in the T4R strain, expression of *SP1434* did not increase following treatment with TPEN in the Δ1433 mutant strain. These data indicate involvement of *SP1433* in the regulation of this operon. To identify if *SP1433* falls within the previously characterized AdcR zinc-dependent regulon, expression of genes known to be regulated by AdcR, *adcA* and *adcAII*, were analyzed in T4R and the Δ1433 mutant. However, no significant differences in expression of *adca* or *adcAII* were detected between the two strains (Supplemental Figure 2), indicating that while the *SP1434-1438* operon is regulated by *SP1433* and is highly responsive to metal-starvation, it is unlikely part of the AdcR regulon and is instead an independent metal sensor.

**Figure 3.**
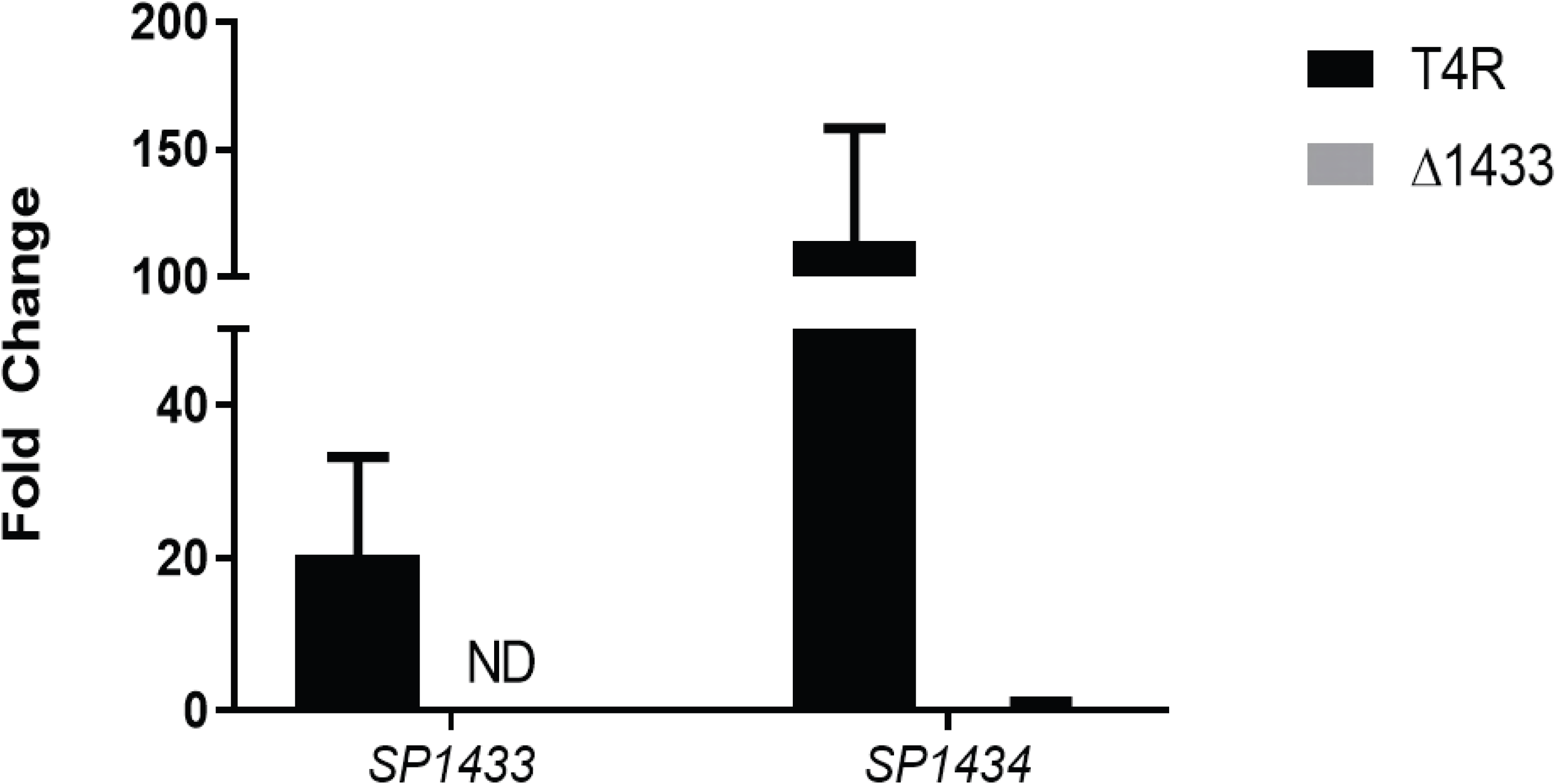
AraC regulator *(SP1433)* regulates operon expression. Gene expression of *SP1433* and *SP1434* were assessed by qRT-PCR in T4R (black) and Δ*SP1433* (gray) strain following 15 min treatment with TPEN (30 μM). Fold changes were calculated by ΔΔCT analysis with *gyrA* serving as an internal control. Non-detectable gene expression is represented by ND.

#### SP1434-1438 Alters Intracellular Metal Availability

The considerable upregulation of this operon following zinc chelation led us to investigate the intracellular metal ion concentrations within T4R and the Δ1434-8 strain using inductively coupled plasma mass spectrometry (ICP-MS). These analyses revealed significant differences in intracellular manganese(II), iron(II), and zinc(II) concentrations, between the two strains (Fig. 4, Supplemental Table 4). Due to sequence similarity with a cobalt/nickel ECF transport system, the difference in intracellular nickel(II) concentration is interesting to note though this was not statistically significant under the conditions tested.

**Figure 4.**
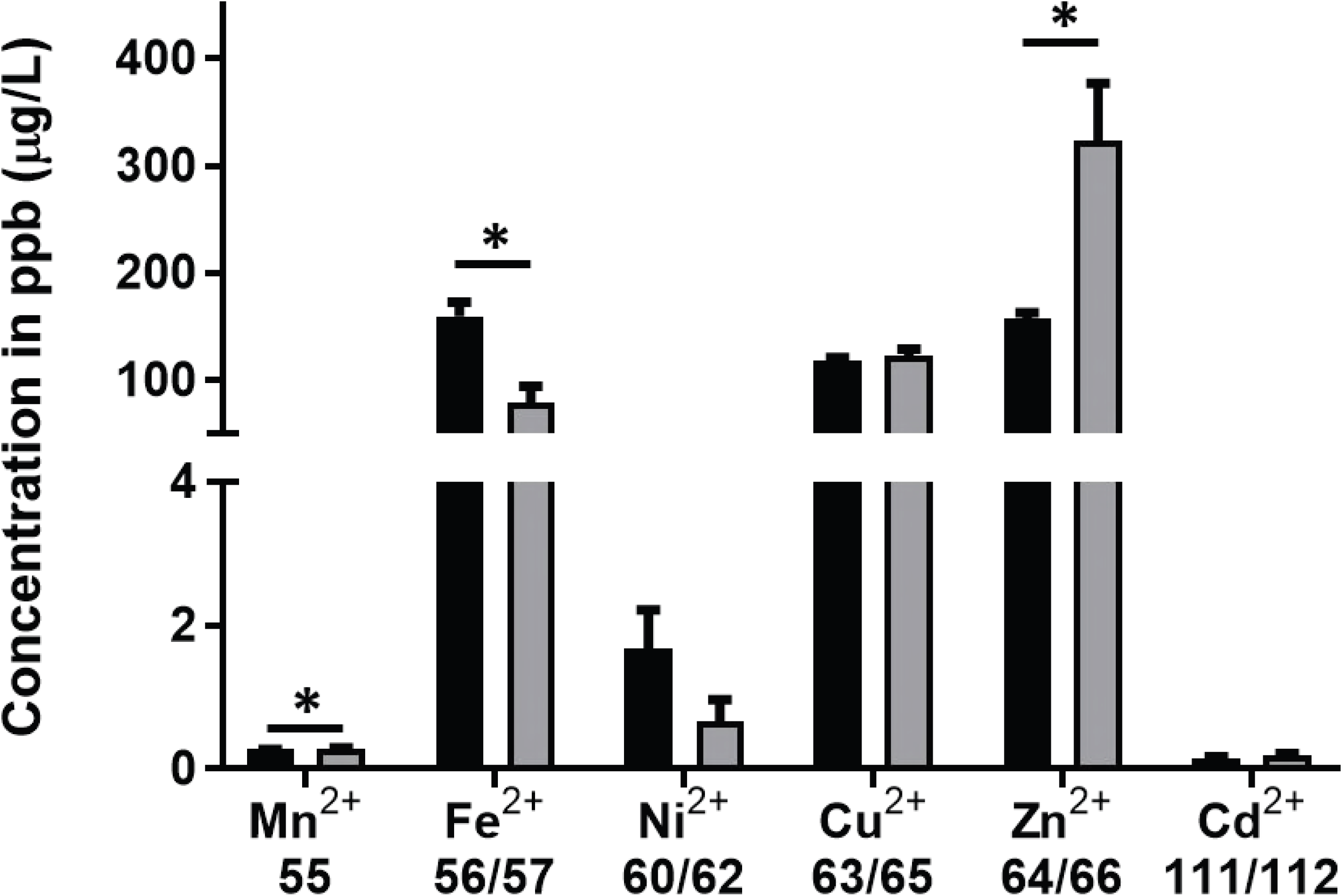
Intracellular metals of T4R and Δ1434-8. a) Metal content of T4R (black) and Δ1434-8 (gray) were assessed by inductively coupled plasma mass spectrometry (ICP-MS) and displayed as concentration in parts per billion (μg/L). Representative figure of three replicates, with bars indicating mean metal concentrations, and standard error of the mean represented by horizontal bars. *p <0.05 as determined by students t-test comparing T4R to Δ1434-8 strain for each individual metal analyzed.

#### Secreted Metabolome of T4R and Δ1434-8

Metals have been shown to impact bacterial metabolism, and a recent study of *Streptococcus pyogenes* showed that excess zinc ions interfere in glucose metabolism through the inhibition of two enzymes: phosphofructokinase and glyceraldehyde-3-phosphate-dehydrogenase (20). To determine the role that the *SP1434-1438* operon is playing in metabolism, potentially due to intracellular metal accumulation, secreted metabolomics were performed on both T4R and Δ1434-8 using a novel 2D nuclear magnetic resonance (NMR) approach. Briefly, cultures of each strain were grown to early log (OD_600_ 0.2), mid-log (OD_600_ 0.35) and beginning stationary phase (OD_600_ 0.5). Culture supernatants were sterile filtered, processed, and quantified by NMR, using a library containing more than fifty metabolites. Partial Least Squares Discriminant Analysis (PLS-DA) of these samples detected significant differences in metabolite concentrations both between strains of pneumococci and across the different optical densities analyzed (Fig. 5A). In addition to identifying the collective differences between strains, significant differences were also detected in individual metabolites between strains, including lactate, acetate, and carbohydrates (Fig. 5B); however, differences were also detected in numerous amino acids including threonine, arginine, and cysteine. A heatmap identifying differences between the two strains throughout the time course of the experiment and clustering metabolites with similar behavior is shown in Supplemental Figure 3. Broadly, the metabolites fall into two main classes: compounds that rapidly increase as OD_600_ increases, including L-cystine, threonine, and lactic acid, and compounds which decrease over time. Additionally, metabolic pathways were mapped showing the most significantly altered metabolites of both T4R and Δ1434-8, with the most metabolic changes occurring during pyruvate metabolism (Fig. 5C). Collectively, these data indicate involvement of this operon in the regulation of cellular metabolism, particularly in glucose and amino acid metabolism.

**Figure 5.**
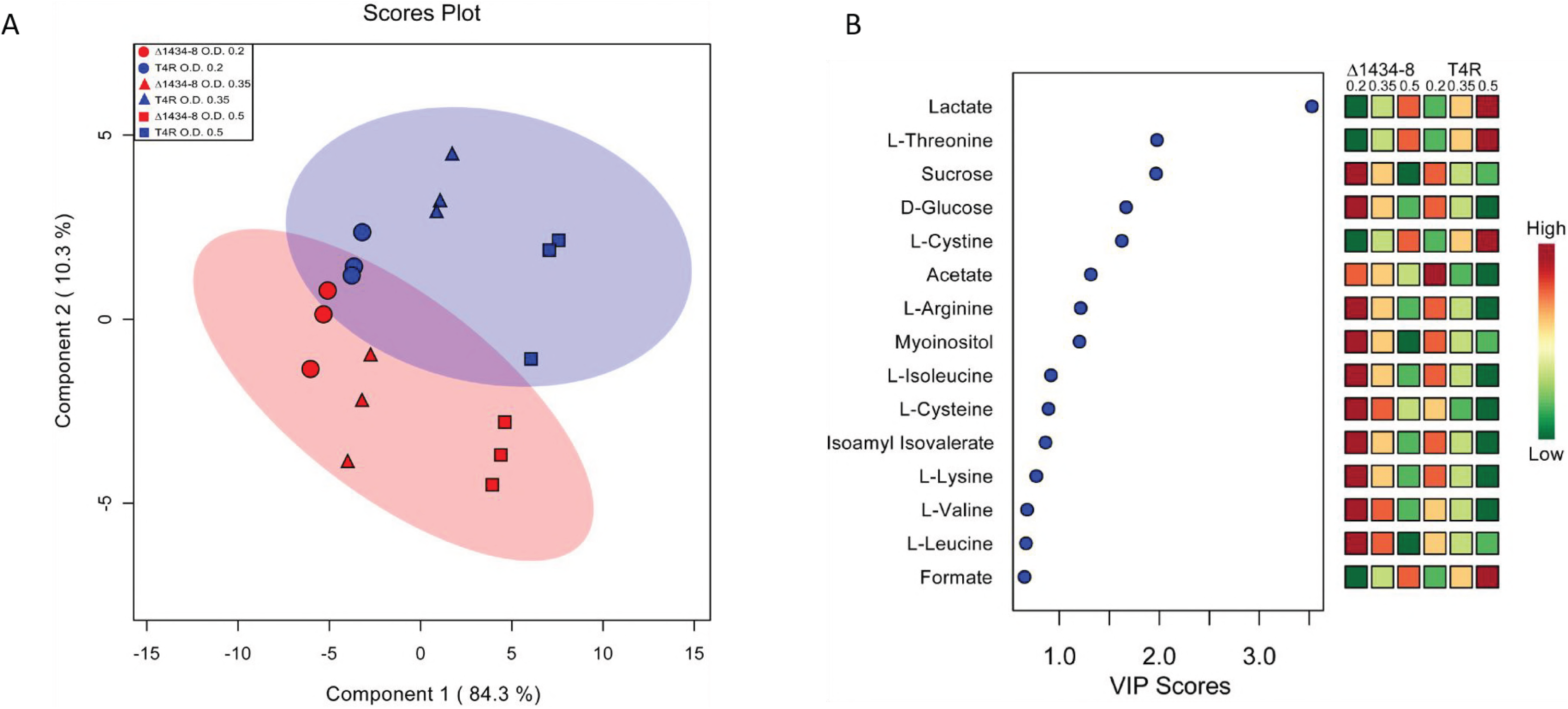
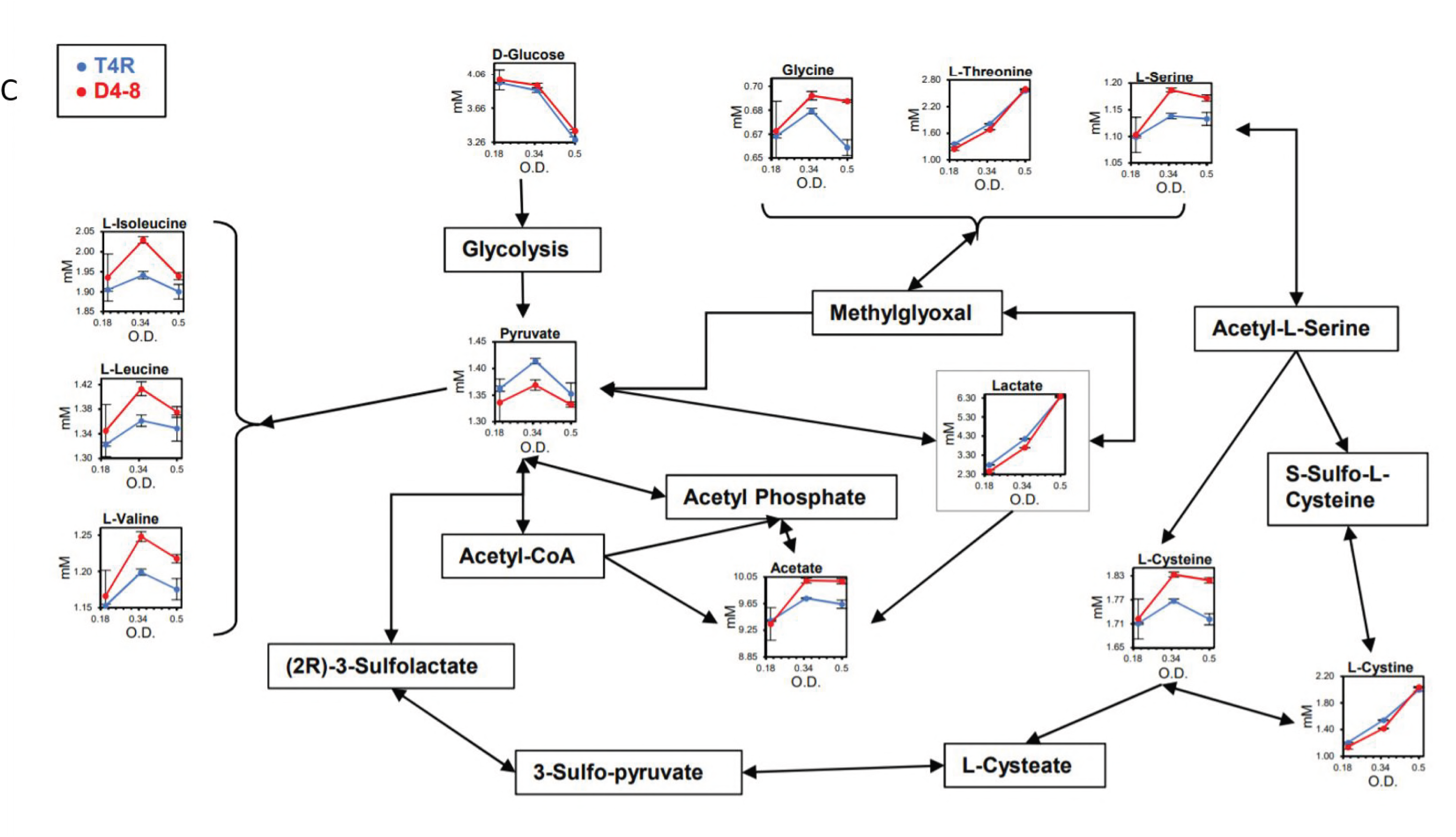
Metabolic profile of T4R and Δ1434-8. A) Clustering of samples within the PLS-DA plot of T4R (blue) and Δ1434-8 (red) profiles indicate significant metabolic differences between the two strains at an OD_600 nm_ of 0.2, 0.35, and 0.5. B) PLS-DA VIP Scores plot of T4R and Δ1434-8 indicates the most significant metabolites identified between strains. C) Extracellular metabolite concentrations of T4R (blue) and Δ1434-8 (red) arranged in metabolic pathways.

#### Proteomic Analysis of T4R and Δ1434-8

To identify potential mechanisms leading to differences in the metabolic profiles between T4R and Δ1434-8 strains, proteomic analysis of both T4R and Δ1434-8 were performed using mass spectrometry-based methods. Using a Fisher’s exact t-test (p < 0.003), we identified 41-differentially expressed proteins in the Δ1434-8 strain compared to the parental T4R (Table 1). Differentially expressed proteins were analyzed through KEGG, STRING, Uniprot, and RegPrecise to determine potentially relevant metabolic and regulatory pathways (21-24). From these data, roughly 22% of those differentially expressed, fall within the CcpA, global catabolite repression regulon (25, 26). Two of the 41 differentially expressed proteins fall under the regulation of CodY, which is also a known global nutritional regulator (27). In addition to the CcpA and CodY regulons, multiple differentially expressed proteins were identified as belonging to the ArgR, the predicted-Rex, and the CtsR regulons, indicating changes in arginine metabolism and redox stimuli between the Δ1434-8 strain and parental T4R (28-31). Furthermore, expression of ABC-transporters or phosphotransferase systems accounted for more than 20% of the downregulated proteins in the Δ1434-8 strain.

**Table 1.**
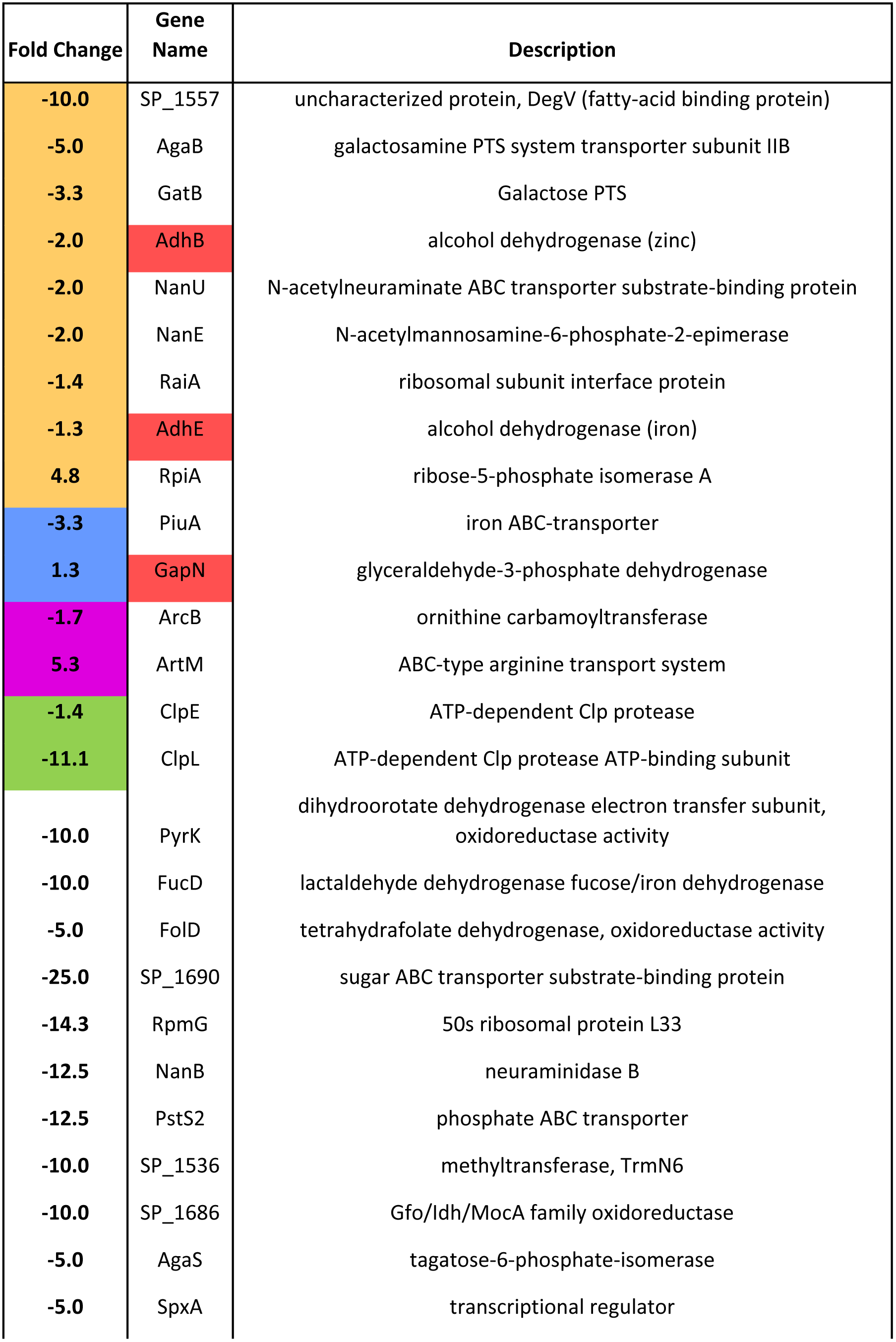
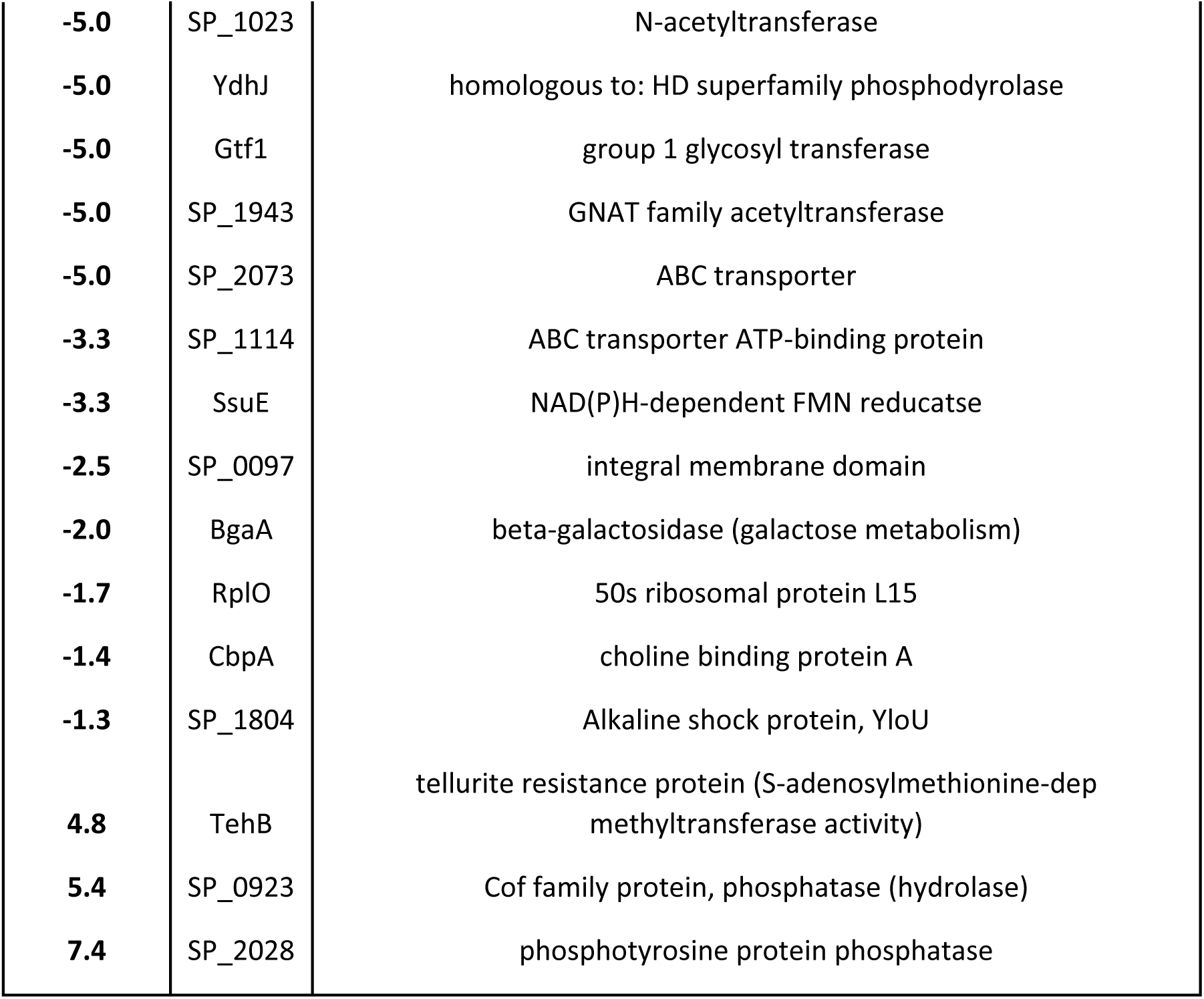
Differentially expressed proteins in ΔSP1434-8 versus T4R. Mass spectrometry based proteomic analysis of Δ1434-8 strain compared to the parental T4R from cultures grown to OD_600 nm_ 0.5 identified 41 -differentially expressed proteins. Fisher’s exact t-test p<0.00274. Highlighted colors represent regulons the proteins are fall within: CcpA (yellow), CodY (blue), Rex (red), CtsR (green), and ArgR (purple).

## Discussion

*Streptococcus pneumoniae* remains one of the leading killers of children worldwide, and infections due to this organism account for more than 400,000 hospitalizations per year in the United States alone (32). The pneumococcus primarily resides as a commensal of the nasopharynx, where zinc(II) concentrations are limited. Additionally, exposure to calprotectin, a protein produced by neutrophils to sequester zinc ions from bacteria, further limits metal availability within the host. Pathogens must therefore utilize mechanisms to circumvent metal starvation. Though zinc(II) acquisition and regulation have been well characterized in S. *pneumoniae*, in this study we have identified a previously uncharacterized system that is strongly responsive to zinc-limitation. Furthermore, loss of this genetic locus results in an altered cellular metabolism. Our results indicate that genes *SP1433-1438* are encoded as an operon that is highly upregulated in response to zinc-chelation by the cell permeable chelator, TPEN, potentially mimicking the environment encountered during neutrophil clearance in the human nasopharynx. BLASTp and UniProt analysis revealed the potential of this operon to encode two transport systems sharing homology to an antibiotic transport system and a cobalt(II)/nickel(II) energy coupling factor transport system. *SP1434* and *SP1435* of this system were recently characterized to be involved in pneumococcal environmental information processing and were found to be highly responsive following deletion of the two-component regulatory system 08 (TCS08) histidine kinase (33). Moreover, the pneumococcal TCS08 has shown to be homologous to the SaeRS system of *Staphylococcus aureus* that is activated by calprotectin, further suggesting the involvement of this system to metal limitation (34, 35).

Due to the promiscuous metal-binding affinity of TPEN, we investigated the effect of other metals on expression of *SP1434*. This operon was determined to be responsive to zinc, cobalt, and nickel ions, as supplementation following TPEN treatment led to a drastic limitation in the upregulation of *SP1434* seen in TPEN treated samples alone. Additionally, we detected upregulation of *SP1434* in the parental T4R strain but no increase in expression in the strain lacking the AraC transcriptional regulator (Δ1433), indicating *SP1433* as the regulator of this operon. Expression of AdcR-regulon genes *adcA* and *adcAII* encoding zinc-binding lipoproteins were not affected in the Δ1433 strain, suggesting that *SP1433* is likely a zinc-sensing regulator functioning independently of AdcR.

Since components of this operon share sequence similarity with metal ion transport systems, we analyzed intracellular metal concentrations of T4R and Δ1434-8 by intercoupled plasma-mass spectrometry (ICP-MS). Significant differences were detected in the concentrations of manganese, zinc, and iron ions. Iron(II) is known to interact with H2O2 through Fenton chemistry to form highly reactive hydroxyl radicals (36), whereas zinc and manganese are known to function as antioxidant metal ions specifically through redox-active metal antagonism and interaction with the superoxide dismutase, SodA (37, 38). As metal ions are also known to be important enzymatic cofactors for proteins involved in metabolism, and H_2_O_2_ is a significant byproduct of pneumococcal metabolism, we explored how loss of this operon, and subsequent changes in intracellular metal ion concentrations, altered cellular metabolism. NMR-based metabolomics of culture supernatants identified major shifts in the metabolic profiles between strains, primarily with increased production of lactate in the T4R strain and increased production of acetate in the Δ1434-8 strain. These differences suggest a preference for mixed acid fermentation by Δ1434-8 versus homolactic acid fermentation by the T4R strain (39). Several amino acids were also observed to exhibit differing metabolite behavior. Of particular interest were the high levels of cysteine detected in Δ1434-8 strain since this strain contains higher concentrations of zinc(II) and cysteine residues are known to interact with zinc ions with high affinity (40).

Proteomic analysis validated the results obtained from the secreted metabolomics and indicated that nearly one quarter of differentially expressed proteins were regulated by the CcpA regulon, involved in carbohydrate catabolite repression. The CcpA regulon has been previously characterized as a master regulator that controls fermentation, as well as catabolism of glucose, galactose, and tagatose (26). Many of the differentially expressed proteins that fall within the CcpA regulon were downregulated and are known to be involved in galactose fermentation, including GatB and the *nan* operon (41, 42). Additionally, CcpA regulation is thought to be important for host interactions and contributes to successful colonization (26, 43). Previous studies have linked sialic acid and CcpA with galactose metabolism specifically through the pneumococcal *nan* operons(42), and it is therefore interesting to note that 4/5 of the genes within this operon in the D39 background (*spd_1263, spd_1264, spd_1265*, and *spd_1267*) were found to be upregulated when grown with 0.5% sialic acid) (42).

In addition to the large number of genes that were identified within the CcpA regulon, results from our proteomic analysis also detected differential expression of proteins within the CodY, ArgR, and Rex regulons. CodY has been shown to be involved in the regulation of colonization and amino acid metabolism and could potentially drive the amino acid differences observed in our metabolomic analyses (44). Arginine metabolism has been linked to virulence in *Streptococcus pneumoniae* and other significant human pathogens and it is interesting to note that one of the six proteins detected to be upregulated in our proteomic analysis was an arginine transport system that is regulated by the ArgR regulon (28). The Rex (redox-sensing regulator) regulon detected by RegPrecise has been characterized in both *Streptococcus mutans* and *Staphylococcus aureus*, though to our knowledge has not yet been characterized in *S. pneumoniae* (45, 46). In both *S. mutans* and *S. aureus* Rex has been shown to sense NAD^+^ or NADH, and in *S. aureus* Rex is thought to be the central regulator of anaerobic metabolism (46). Findings from our proteomics analyses indicate that the involvement of Rex in this system is at the level of a zinc(II) and iron(II)-binding alcohol dehydrogenase, both of which are also under the regulation of CcpA. An additional protein thought to be under coregulation by Rex and CodY that was identified in our proteomics data is a glyceraldehyde-3-phosphate-dehydrogenase. Collectively, these data indicate major metabolic differences between our T4R strain and the Δ1434-8 strain, particularly in carbohydrate metabolism, fermentation, and amino acid metabolism.

In this study, we identified a previously uncharacterized operon of *Streptococcus pneumoniae* that is strongly responsive to zinc-chelation, yet independent of the AdcR zinc(II) regulon. We have identified the regulator for this genetic locus and have determined that mutants lacking the operon (Δ1434-8) display different intracellular metal ion ratios and altered metabolic profiles. Analysis of the secreted metabolomes and proteomic profiles suggest changes in central carbohydrate metabolism, potentially through a shift in fermentation pathways. These data demonstrate that the metabolome of *Streptococcus pneumoniae* is largely metal-dependent, which to our knowledge has not yet been characterized. This work provides a foundation for identifying key metabolic enzymes and intermediates that could be targeted using metal-dependent therapeutics.

## Materials and Methods

### DNA Manipulation

*S. pneumoniae* strains TIGR4 and its unencapsulated mutant (T4R) were grown on tryptic soy agar plates supplemented with 5% defibrinated sheep blood or in C+Y medium. Mutants of TIGR4 and T4R lacking *SP1434-1438* (Δ1434-8) and SP1433 (Δ1433) were created using splicing by overlap extension (SOE) PCR method using an erythromycin or spectinomycin antibiotic cassette and standard *S. pneumoniae* transformation procedures. Mutants lacking SP1434-1438 and SP1433 were isolated by selection on blood agar plates supplemented with erythromycin (0.5 μg/mL) or spectinomycin (500 μg/mL), respectively, and confirmed by PCR.

### Antibiotic Sensitivity

Frozen bacterial stocks of T4R and Δ1434-8 were diluted to 1×10^7^ CFU/mL and 100 μL were spread on a blood agar plate. Discs impregnated with antibiotics at the following amounts were added to blood agar plates and were incubated overnight at 37 °C with 5% CO_2_ (Ciprofloxacin 5 μg, Vancomycin 30 μg, Ampicillin 10 μg, Penicillin 10 U, Ceftiofur 30 μg, Cephalothin 30 μg, and Sulfisoxazole 1 mg). Following incubation, zones of inhibition surrounding antibiotic discs were measured.

### Inductively Coupled Plasma Mass Spectrometry

Bacterial cultures of T4R and Δ1434-8 were grown to OD_600_ nm 0.6 in triplicate. Four 1 mL cultures were collected of each strain, centrifuged at 16,000xg for 5 min, and supernatant was decanted. Pellets were heat killed at 65 °C for 2 hr. Pellets were resuspended in 100 μL concentrated nitric acid, then were diluted 1:20 with water. An Agilent ICP-MS 7500cx was used to collect all ICPMS data herein.

### Quantitative Real-time PCR

For assays investigating *SP1434* expression, *S. pneumoniae* T4R was grown in C+Y medium to O.D._600 nm_ of 0.6 prior to the addition of the Zn^2+^-chelating agent TPEN (30 μM) or zinc(II), cobalt(II), iron(II), or nickel(II) ions at 200 μM. Supplemented metals were TraceCERT^®^ ICP-MS grade (Sigma-Aldrich). After addition of TPEN or individual metals, bacteria were incubated at 25°C for 15 min. Additionally, cultures were exposed to TPEN for 15 min followed by metal for a following 15 min. For assays investigating *SP1433* expression, T4R and Δ1433 were grown to O.D._600 nm_ of 0.6 and treated for 15 min with TPEN (30 μM). For all samples, after incubation, 1 mL of bacterial culture was added to 2 mL of RNAprotect (Qiagen). Samples were incubated at room temperature for 5 min. Two mL of bacterial culture in RNAprotect was pelleted for 5 min at 16,000×g, pellets were resuspended in 1 mL of cold RNase free PBS and centrifuged again. Supernatants were decanted, and pellets were resuspended in 400 μL RLT buffer (Qiagen) with 2-mercaptoethanol (Sigma-Aldrich). Samples were sonicated three times (15 sec), 600 μL RLT buffer was added to each sample, and samples were transferred to 500 μL of 0.7 mM Zirconia beads. Samples were bead beat for 2 min using Mini-BeadBeater 16 (BioSpec Products). Lysates were centrifuged on tabletop centrifuge at 2,000xg for 1 min. Samples were run through Qiashredder columns (Qiagen) per manufacturer’s instructions. 100% Ethanol was added to Qiashredder flow through at 0.6 volume of the sample. RNA was purified using a Qiagen RNeasy Mini Kit (Qiagen), optional on-column DNase treatment was performed for 30 min, RNA was then quantitated using a Qubit, and 5 ng of each sample was used to synthesize cDNA using a Maxima First Strand cDNA synthesis kit (Thermo Scientific). cDNA products were diluted 1:10 and 1 μL of cDNA was used as template for qRT-PCR using Luminaris Color HiGreen High ROX qPCR Master Mix (Thermo Scientific) per manufacturer’s instructions. Primer sequences can be found in Supplemental Table 3.

### Secreted Metabolomics

Cultures of T4R and Δ1434-8 were grown to OD_600 nm_ 0.2, 0.35 and 0.5. One milliliter of culture was removed and centrifuged at 16,000×g for 5 min and sampling was performed in triplicate. Culture supernatants were then filtered using 0.22 μM filters. NMR samples were made by combining the filtered supernatant (400 μL) with 200 μL of 200 mM phosphate buffer (pH 7.0) with 1.000 mM trimethylsilypropanoic acid (TMSP) in 50% D_2_O. The 1D and 2D NMR spectra were obtained at a temperature of 298 K on a 600 MHz Bruker Avance III cryoprobe equipped NMR spectrometer. A 1D-NOESY (noesypr1d) pulse sequence was used, and presaturation applied at 4.75 ppm during the 4 second relaxation delay and 50 millisecond mixing time to suppress the water signal. A 1 second acquisition time was used, and a total of 64 scans were collected with a 20-ppm spectral width. A modified 2D-TOCSY (dipsi2gpphzsprespe_psyche) pulse sequence was used for 2D acquisition. This sequence incorporated broadband homonuclear decoupling using PSYCHE in the *t_1_* dimension(47-49). A zero-quantum filter was used during the 80 ms TOCSY mixing time. Water flip back pulses were used to optimize water suppression (50). Each FID was collected using 2 scans, and the indirect dimension was sampled for 85 ms using 1024 complex points. Additional water suppression and solvent filtering were performed with NMRPipe using in-house scripts (51). Each spectrum was calibrated using the TMSP peak as an internal standard and manually processed. Compounds in the processed spectra were identified and quantified using AMIX-Viewer v3.9.14 software (Bruker Biospin GmbH). A library of 56 pure compounds at 3.000 mM was created in-house to identify and quantify peak intensities and line widths for each compound. The output file from AMIX was a listing of concentration for each compound for each sample. Concentrations for each sample (T4R and Δ 1434-8) and O.D. (0.2, 0.35 and 0.5) were used for the statistical analysis.

The metabolite concentrations for each sample was arranged by O.D. values and strain before the statistical analysis was conducted using MetaboAnalyst(52). MetaboAnalyst normalized the samples by sum with Pareto scaling. The Pareto scaling was used to emphasize the weaker metabolites and reduce the influence of the intense peaks to easily identify the biological relevance(53). After normalization, the statistical methods, such as multivariate analysis, were used for data analysis. The PLS-DA data set was divided into components to identify the statistical differences between the classes. The first component (Component 1) captured the maximum variance in the data set that was the linear combination of the original predictor variables compared to the observed variables, whereas the other components (second, third, fourth, etc.) captured the remaining variance in the data set that was the linear combination and orthogonal to the first component(53).

### Proteomic Analysis

Four cultures of T4R and Δ1434-8 were grown to OD_600 nm_ 0.4 in C+Y medium and subjected to liquid chromatography-tandem mass spectrometry (LC-MS/MS) analysis as previously described (54, 55). Briefly, proteins were isolated from bacterial pellets sonicated in NP-40 lysis buffer (0.5% NP-40, 150 mM NaCl, 20 mM CaCl2·2H_2_O, 50 mM Tris, pH 7.4) supplemented with protease inhibitor cocktail (c0mplete™, Sigma-Aldrich) using a Covaris S220 focused-ultrasonicator. Protein concentration was determined using Thermo Scientific Pierce BCA Protein Assay Kit. Precipitation of 30 μg of protein was performed with methanol and chloroform (4:1), solubilized in 8 M urea, reduced (5 mM dithiothreitol (DTT) at 65 °C for 10 m) and alkylated (0.01 M iodoacetamide at 37 °C for 30 m) and digested with porcine trypsin (at 37 °C, overnight, 50:1 ratio of protein. Tryptic peptides were desalted using a C18 spin column (Thermo Fisher Scientific) and analyzed by linear trap quadropole (LTQ) Orbitrap Velos mass spectrometer equipped with an Advion nanomate electrospray ionization (ESI) source (Advion). Peptides (500 ng) were eluted from a C18 column (100 μm id × 2 cm) onto an analytical column (75 μm ID × 10 cm, C18) using a 180 m gradient with 99.9% acetonitrile, 0.1% formate at a flow rate of 400 nL/m and introduced into an LTQ-Orbitrap.

Data dependent scanning was performed by the Xcalibur v 2.1.0 software using a survey mass scan at 60,000 resolution in the Orbitrap analyzer scanning mass/charge (m/z) 400–1600 followed by collision-induced dissociation (CID) tandem mass spectrometry (MS/MS) of the 14 most intense ions in the linear ion trap analyzer (56). Precursor ions were selected by the monoisotopic precursor selection (MIPS) setting with selection or rejection of ions held to a ±10 ppm window. Dynamic exclusion was set to place any selected m/z on an exclusion list for 45 s after a single MS/MS. Tandem mass spectra were searched against a *Streptococcus pneumoniae* serotype 4 strain ATCC BAA/TIGR4 FASTA protein database downloaded from UniProtKB to which common contaminant proteins (e.g., human keratins obtained at ftp://ftp.thegpm.org/fasta/cRAP) were appended. All MS/MS spectra were searched using Thermo Proteome Discoverer 1.3 (Thermo Fisher Scientific) considering fully tryptic peptides with up to two missed cleavage sites. Variable modifications considered during the search included methionine oxidation (15.995 Da), and cysteine carbamidomethylation (57.021 Da). Peptides were identified at 99% confidence with XCorr score cutoffs based on a reversed database search (57). The protein and peptide identification results were visualized with Scaffold v 3.6.1 (Proteome Software Inc.). Protein identifications with a minimum of two peptides identified at 0.1% peptide false discovery rate (FDR) were deemed correct. Significant changes in protein expression between T4R and Δ1434-8 were identified by Fisher’s exact test at a p-value of ≤0.054 and fold change of ±1.3. Fold changes in protein expression were calculated using weighted normalized spectra with 0.5 imputation value. Various bioinformatics resources such as DAVID, KEGG, and STRING were utilized to determine the functions of the identified proteins (22, 58, 59). The PRoteomics IDEntifications (PRIDE) database is a centralized, standards compliant, public data repository for proteomics data. The mass spectrometry proteomics data from this study is deposited to the ProteomeXchange Consortium via the PRIDE partner repository (pending accession number) (60).

### Statistics

Intercoupled plasma mass spectrometry (ICP-MS), and H_2_O_2_ killing assays were performed a minimum of three times, results from independent experiments were averaged together, and standard error of the mean was calculated. Data sets were analyzed by comparing parental T4R to Δ1434-8 using the students t-test, with an α value = 0.05. Results were deemed statistically significant when p < alpha. Statistical analyses were performed using GraphPad Prism 7.

## Acknowledgements

Thank you to Allen Shack in the College of Basic Sciences and the staff of the Arizona State University Mass Spectrometry Core for their assistance with the proteomic analysis featured in the work presented here. This work was supported in part an Institutional Development Award (IDeA) from the NIGMS COBRE grant number P20GM103646 (awarded to JAT) and by a National Institutes of Health grant number R15GM113152 (awarded to NCF).

**Supplemental Table1.** RNA was harvested from bacterial cultures of TIGR4 and TIGR4 treated with TPEN grown to OD _600 nm_ 0.5 and was used to synthesize cDNA for hybridization to pneumococcal microarray.

**Supplemental Table2.** Blast analysis of S. *pneumoniae* TIGR4 proteins SP1433-SP1438 revealed homology to a transcriptional regulator and two transport systems.

**Supplemental Table3.** Primers sequences used in this study for molecular cloning and gene expression studies by qRT-PCR.

**Supplemental Figure 1.** Antibiotic sensitivity of T4R and ΔSP1434-8. T4R (black) and ΔSP1434-8 (gray) were inoculated onto blood agar plates and antibiotic impregnated discs were added at the following concentrations: Ciprofloxacin 5 μg, Vancomycin 30 μg, Ampicillin 10 μg, Penicillin 10 U, Ceftiofur 30 μg, Cephalothin 30 μg, and Sulfisoxazole 1 mg. Following overnight incubation, zones of inhibition surrounding antibiotic discs were measured (mM).

**Supplemental Figure 2.** SP1433 specifically regulates SP1434-8. Gene expression of *adcAII, adcA, SP1434* and *SP1433* were assessed by qRT-PCR in T4R (black) and *ASP1433* (gray) strain following 15 min treatment with TPEN (30 μM). Fold changes were calculated by ΔΔCT analysis with *gyrA* serving as an internal control. Non-detectable gene expression is represented by ND.

**Supplemental Figure 3.** Heatmap of secreted 2D-NMR metabolomics. Cultures of T4R and Δ1434-8 were grown to OD_600 nm_ 0.2, 0.35, 0.5. Supernatants were collected, sterile filtered, and analyzed against a metabolite library.

**Supplemental Table4.** Mean intracellular metal ion concentrations in ppb (μg/L) in both the T4R and Δ1434-8 strains with standard deviation shown in parenthesis.

